# Systematic auditing is essential to debiasing machine learning in biology

**DOI:** 10.1101/2020.05.08.085183

**Authors:** Fatma-Elzahraa Eid, Haitham Elmarakeby, Yujia Alina Chan, Nadine Fornelos Martins, Mahmoud ElHefnawi, Eli Van Allen, Lenwood S. Heath, Kasper Lage

## Abstract

Representational biases that are common in biological data can inflate prediction performance and confound our understanding of how and what machine learning (ML) models learn from large complicated datasets. However, auditing for these biases is not a common practice in ML in the life sciences. Here, we devise a systematic auditing framework and harness it to audit three different ML applications of significant therapeutic interest: prediction frameworks of protein-protein interactions, drug-target bioactivity, and MHC-peptide binding. Through this, we identify unrecognized biases that hinder the ML process and result in low model generalizability. Ultimately, we show that, when there is insufficient signal in the training data, ML models are likely to learn primarily from representational biases.

---

Datasets in the life sciences have grown increasingly large and complicated. With the advent of single cell studies and biobanks, scientists are turning to machine learning (ML) to derive meaningful interpretations of these massive genomic, transcriptomic, proteomic, phenotypic, and clinical datasets. One major obstacle to the development of reliable and generalizable ML models is that auditing for biases has not been established as a common practice in life sciences ML; despite a large body of work in non-biological ML which addresses the identification and removal of algorithm biases (Zou and Schiebinger 2018). Yet, it is well known that biological datasets often suffer from representational biases stemming from evolutionary, inherent, and experimental artifacts. When these biases are not identified and eliminated, the ML process can be misled such that the model learns predominantly from the biases unique to the training dataset and, hence, is not generalizable across different datasets. In this scenario, prediction performance is inflated for the test set, but drops drastically for external predictions. Here, we demonstrate that several prominent protein-protein interaction (PPI) predictors are overwhelmingly reliant on a bias in their training dataset - to the extent that the PPI predictions become randomized (i.e., the model has no predictive power) once the bias is removed. For this reason, when applying ML to biological datasets, it is crucial to systematically audit for biases inherent in the data. This will help us to understand how and what the model is learning in order to ensure that its predictions are based on true biological insights from the data.

We devised a systematic auditing framework for paired-input biological ML applications (**Fig. 1a**), which are widely harnessed to predict the biological relationships between two entities, e.g., physical interactions between proteins, bioactivity of drugs and their targets, or binding of MHC molecules to their antigens. We used this framework to identify biases that have confounded, over the past two decades, the ML process in three applications of great interest to the life sciences and biotech communities: protein-protein interactions, drug-target bioactivity, and MHC-peptide binding.

**Figure 1.**
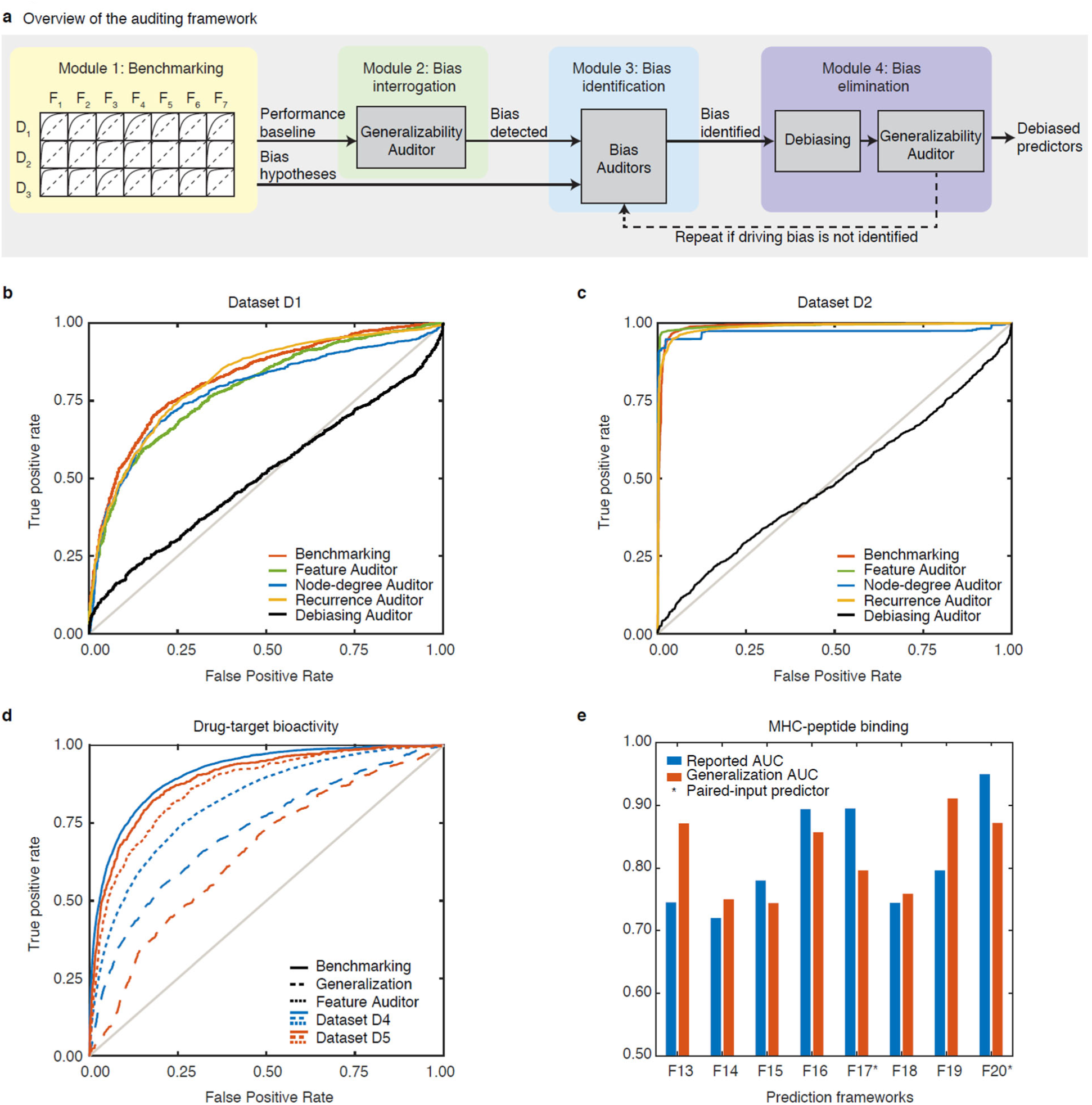
A systematic auditing framework for ML applications in biology. (**a**) The four modules of the auditing framework. (**b**) The performance of the best performing PPI classifier, F1, on dataset D1 for benchmarking and the four auditors. In the Feature Auditor, protein features are masked; yet, classifier performance is retained. In the Node-degree and Recurrence Auditors, proteins are represented solely by their node degrees in the positive or negative training dataset or by their differential node degree between the positive and negative training examples; yet, without protein features, classifier performance informed by node degree or protein recurrence alone is retained. (**c**) The performance of the best performing PPI classifier, F5, on dataset D2, with similar observations as with **Fig. 1b**. (**d**) The performance of the optimized drug-target predictor F8 on datasets D4 and D5 in the benchmarking (module 1), generalization (module 2), and bias identification (module 3, *Feature Auditor*). (**e**) The reported (module 1) and generalization (module 2) performances of MHC-peptide binding predictors F13-F20. Asterisks denote the two paired-input predictors, F17 and F20.

## Protein-protein interaction predictors

Mapping protein-protein interactions (PPIs) is critical to understanding cellular processes, interpreting genetic data, and predicting new targets for therapeutics development. This has led to a great interest in developing PPI classifiers that are designed to learn from interacting (positive) and non-interacting (negative) protein pairs to determine which protein features (*e.g.*, amino acid physicochemical properties) contribute to an interaction between two proteins. In particular, the holy grail of PPI classifiers is the ability to predict PPIs based on nothing but protein sequence - with no input describing structure or evolutionary information. There are existing models that learn from protein domain structure to predict interactions (Lasso et al. 2019). However, these are limited by the requirement for PPI structure characterization - these are uncommon, especially for novel proteins that are the key targets of PPI predictors; we typically endeavor to predict interactions for proteins that are not characterized rather than proteins for which structural data exist - and cannot be easily extrapolated to peptides that are structurally flexible. These limitations drive the demand for PPI predictors that rely on amino acid biochemical and physical properties alone. At the very least, PPI classifiers are supposed to acquire the ability to infer whether a given protein pair is likely to interact based on protein features, rather than biases that plague a given dataset.

A critical and unexplained observation regarding PPI classifiers is that they achieve very high and, sometimes, near-perfect performances (Pan, Zhang, and Shen 2010; Shen et al. 2007; Sun et al. 2017; Park and Marcotte 2012). This is despite the fact that the models use simple features such as protein amino acid sequence, which, from a biochemical and molecular biology perspective, should not be sufficient to determine physical interactions between proteins. Specifically, the feature designs do not take into consideration which protein residues contribute to interactions, the spatial relations among residues, or even the location of the protein within cellular compartments. A central question in the field is therefore, what PPI classifiers are learning from such simple protein sequence features in order to almost perfectly predict PPIs. We hypothesized that unidentified biases in the training data may be driving the high performance. To test this hypothesis, we devised an auditing framework composed of four main modules: benchmarking, bias interrogation, bias identification, and bias elimination (**Fig. 1a**, and **Online Methods**).

In the first benchmarking module, we benchmarked classifiers on different datasets to establish a baseline performance for subsequent comparisons and to identify performance patterns suggestive of data biases. We selected seven prominent PPI classifiers, which we refer to as F1-F7 in this work, each represented by a variety of ML algorithms (support vector machines, random forests, and deep neural networks) and characterized by diverse protein feature descriptors (e.g., frequency of k-mers, autocorrelation between amino acid properties, and domain occurrences). The performances of F1-F7 were benchmarked on two curated PPI datasets, D1 (Park and Marcotte 2012) and D2 (Pan, Zhang, and Shen 2010), which are widely used to develop and test PPI classifiers; and D3 (Rolland et al. 2014), a high-quality experimental dataset. All three datasets involve human proteins and are highly relevant for the development of therapeutics. Classifiers were trained on subsets of a specific dataset and tested on non-overlapping subsets of the same dataset. As anticipated, the best **benchmarking** performance across all classifiers was high with an average area under the curve (AUC) of 0.83, 0.99, and 0.92 for D1, D2, and D3, respectively (**Supplementary Table 1**). The performances we measured are similar to the published performances of F1-F7, indicating the correct implementation of the classifiers.

In the second module, we examined biases in the training data by assessing the ability of each classifier to generalize to independent datasets. Briefly, before we expand on this module, there are two similar ideas that exist in the biological ML community that need to be discussed first. One, it has been posited that the bias of a training dataset is valuable for prediction-making, e.g., the frequency of a given protein in the positive versus the negative training data is useful for the training of ML models. In other words, proteins that have many interaction partners are generally more likely to interact with a random given protein, and interactions are much more likely between two proteins that both have many interaction partners (**Supplementary Discussion**). Two, many PPIs are known to be context specific, e.g., tissue- or disease-specific. We would like to counter both points before we demonstrate the importance of removing biases from training datasets. Mainly, although many PPIs are context-specific, a considerable fraction of PPIs are shared across human cells and PPIs are governed by universal principles that ML models are supposed to learn from. If a given predictor’s predictions become entirely random (akin to guessing without any information which proteins interact with each other) after training biases have been removed, this indicates that the predictor was predominantly learning from the bias and is not generalizable. In such cases, the bias is not empowering predictions, but rather lobotomizing the ML process so that it does not discern the true principles that determine protein-protein interaction, even within a given context or network.

Therefore, in this second module, we designed a *Generalizability Auditor* that examines how F1-F7 perform on a dataset independent from that used for training (we used subsets of D1 and D2 for training and subsets of D3 for independent testing). In the absence of bias, the AUCs of the *Generalizability Auditor* and the benchmarking in the first module should be near-identical. In contrast, we observed a significant difference: AUCs of the *Generalizability Auditor* are significantly lower by an average of 0.14 and 0.33 for the classifiers trained on D1 and D2, respectively (**Supplementary Fig. 1**). This suggests that there are dataset-specific biases, which is more pronounced for D2, that confound the learning process and inflate the performances of F1-F7.

In order to determine what these biases are, in the third module, we followed a principled cycle of steps where hypotheses regarding potential biases are iteratively formulated and tested via hypothesis-specific auditors. These auditors are auxiliary ML models, each designed to assess a specific hypothesis about any given ML model of interest, i.e., F1-F7. We started by testing the hypothesis that each PPI classifier is learning solely from protein features, as it is designed to, by conceiving a ***Feature Auditor*** that masks all protein features simultaneously. When the protein features are masked, the auditor performance should become random. Yet, we found that the benchmarking performance of each PPI classifier was largely retained regardless of the underlying ML classifier, hyperparameter values, training dataset, or protein features: average difference in AUC is −0.01, 0.00, and −0.01 for D1, D2, and D3, respectively (**Fig. 1b, 1c**, and **Supplementary Table 1**). This rejects the hypothesis that F1-F7 are learning from protein features.

We next hypothesized that protein recurrence in the training data was inflating the performance. This hypothesis was inspired by a comment by Park & Marcotte that “if an object is present more often in positive than in negative training pairs, most predictive algorithms successfully learn that test pairs involving that object are more likely to interact than not, which often turns out to be true” (Park and Marcotte 2011). We sought to understand the extent to which protein recurrence in the positive versus negative training dataset was dictating predictions. To test this, we built a ***Node-degree Auditor*** in which each protein was solely represented by its node degrees in the positive and negative training examples. The performance using the ***Node-degree Auditor*** was highly similar to the best benchmarking performance across all classifiers for each dataset: difference in AUC is 0.03, 0.01, and 0.05 for D1, D2, and D3, respectively (**Fig. 1b, 1c**). These results confirm the hypothesis that protein recurrence is largely informing and inflating the performance of F1-F7.

Based on the observations from the ***Feature Auditor*** and ***Node-degree Auditor*** - that F1-F7 are not learning from protein features but rather from protein recurrence in the training datasets - we built a ***Recurrence Auditor*** that uses differential node degrees of a given protein between the positive and negative training examples as the sole information to estimate the probability of interactions for new protein pairs. Unfortunately, the performance using the ***Recurrence Auditor*** is similar to the best benchmarking performance, even more so than the ***Node-degree Auditor***, across all classifiers for each dataset: difference in AUC is 0.00, 0.04, and 0.06 for D1, D2, and D3, respectively (**Fig. 1b, 1c**). This confirms that the performance of F1-F7 is primarily determined by the difference in protein recurrence in the positive versus negative training dataset.

Finally, we implemented a ***Debiasing Auditor*** that accounts for protein node degree bias by ensuring that each protein has an equal node degree in the positive and negative training examples while protein features are masked. If node degree is a strong performance driver, this auditor should exhibit near-random performance. As expected, the predictions were effectively randomized across all combinations of classifiers, hyperparameter values, training datasets, and protein features: average AUC is 0.50, 0.53, and 0.49 for D1, D2, and D3, respectively (**Fig. 1b, 1c**, and **Supplementary Table 1**), confirming the hypothesis that node degree is strongly biasing the PPI classifier performance. In other words, F1-F7 are not learning from protein features to predict PPIs; instead these classifiers use the bias inherent in the training datasets to predict PPIs.

In the fourth and final module, we remove the biases identified in the third module and use the *Generalizability Auditor* to assess how the classifiers generalize to independent datasets after debiasing. If we have effectively eliminated biases in the training dataset, the benchmarking performance (**Supplementary Table 1**) and the performance of the *Generalizability Auditor* should be similar. We applied the *Generalizability Auditor* to F1-F7 with debiased training subsets of D1 and D2, distinct benchmarking subsets of D1 and D2, and subsets of D3 for independent testing. As predicted, this improved the PPI classifier generalizability: average difference in AUC between the benchmarking and independent testing performances is 0.06 and 0.03 for D1 and D2, respectively, compared to 0.14 and 0.33 in the first module (**Supplementary Fig. 1**). However, the overall generalizability performance is low, indicating that the PPI predictors still do not learn adequately even when the bias has been removed.

## Drug-Target and MHC-peptide predictors

To illustrate the broad applicability of our auditing framework, we adapted it to two additional applications of significant therapeutic interest: predictions of drug-target bioactivity and MHC-peptide binding. For drug-target bioactivity prediction, we examined five predictors: three classification and two regression frameworks, F8-F12, on two datasets, D4 and D5. Once again, the drug-target predictors did not generalize as well as their benchmarking performance (**Supplementary Table 2** and **Fig. 1d**). Although these predictors are not immune to the protein recurrence bias, they are not impacted to the same extent as the examined PPI predictors. Notably, the biases in each dataset vary: The F8 predictor is less impacted when drug/protein features are masked in D4 as compared to D5; F8 is less generalizable when it learns from D5 compared to D4. This indicates that the dataset D4 has relatively more biological signal and less bias compared to D5 for predictor F8 to learn from. The extent to which a biological dataset is biased could be influenced by numerous factors. For example, alongside the presence of “promiscuous” proteins that bind to many other proteins, the size of the dataset and the experimental assay utilized to collect the dataset can significantly influence bias.

For MHC-peptide predictions, we examined eight predictors, F13-F20. Only F17 and F20 utilize a paired-input setting and exhibit much lower generalized performance compared to reported performance (**Fig. 1e**). The remaining six predictors built separate models for each MHC allele, i.e., a set of single-input models for each predictor instead of a single paired-input model. Overall, drug-target and MHC-peptide predictors generalize in a non-random fashion, suggesting that they learn from their input features in a more biologically meaningful way compared to the examined PPI predictors.

## Conclusion

In conclusion, applying our framework to predictors from three different ML applications suggests that, when there is insufficient signal in the training data, ML models could learn primarily from representational biases or other biases in the training data. This appears to predominantly influence paired-input ML applications, and can be highly misleading if not illuminated through auditing. We recommend that scientists who are applying ML to biological applications help to build a community-wide stance on the systematic auditing of ML models for biases. Being cognizant of the biases that fuel the predictions of each ML model will inform their application to new datasets and clarify whether the model has truly learnt from governing biological principles.

## Acknowledgments

We thank Yu Xia (McGill University), Paul A. Clemons (Broad Institute of MIT and Harvard), and Lucas Janson (Harvard University) for their helpful discussions, and Shuyu Wang (UCSF) for preparation of the localization data. This work was supported by grants from The Stanley Center for Psychiatric Research, the National Institute of Mental Health (R01 MH109903), the Simons Foundation Autism Research Initiative (award 515064), the Lundbeck Foundation (R223-2016-721), a Broad Next10 grant, and a Broad Shark Tank grant. Y.A.C. was funded by a Human Frontier Science Program Postdoctoral Fellowship [LT000168/2015-L].

## Supplementary Materials

### Supplementary Methods

#### Datasets

D1, a curated dataset, contains 24,718 positive protein-protein interaction (PPI) examples among 7,033 human proteins that share at most 40% sequence identity (Park and Marcotte 2012). D1 follows the random negative sampling scheme, which is the most commonly utilized for negative training in PPI classification frameworks: negative PPI examples are generated by randomly pairing proteins not reported to interact in the dataset. D2, another curated dataset, has a predefined pool of negative examples generated by pairing proteins (from the positive example pool) that do not colocalize in the same subcellular compartment. The negative and positive PPI examples in D2 number 36,320 each, among 10,336 human proteins (Pan, Zhang, and Shen 2010; Sun et al. 2017). D3, available at http://interactome.dfci.harvard.edu/H_sapiens/host.php, is a set of 15,473 PPIs among 4,569 human proteins identified using a high-quality all-versus-all Y2H system such that pairs not identified as positive PPIs can be considered experimentally negative (Rolland et al. 2014). Here, each dataset is split into 10 rounds of training, validation, and test sets. Positive and negative examples are of equal count throughout the entire study to avoid class imbalance. The testing is limited to in-network test sets throughout the study, i.e., proteins in the testing sets must have examples of their other interactions in the corresponding training sets because PPI predictors do not generalize to out-of-network predictions where one or the two proteins of a test pair has no examples of their other interactions in the training sets (Park and Marcotte 2012).

One well-appreciated challenge in the development of PPI predictors is the absence of gold standard negative training examples. This is because biological studies typically verify positive PPI examples, but do not determine the absence of interactions between given proteins. D1 sought to eschew this problem by randomly calling PPIs as negative because the majority of proteins are not expected to interact with each other. This approach is the most widely used in PPI prediction data preparation. D2 adopted a different approach by randomly calling PPIs between proteins from different cellular locations as negative because proteins in different locations are not expected to interact with each other generally. D3 considered PPIs not identified in their Y2H screen to be negative examples. However, all three approaches suffer from their own unique caveats. Unlike positive PPI examples, negative PPI examples require much more validation to conclusively determine that a particular pair of proteins do not interact.

#### PPI classification frameworks

The utilized PPI prediction frameworks, F1-F7, all use amino acid sequences as their sole source of protein features but vary in their feature designs and machine learning (ML) models. They were selected based on their reported high performance, popularity, and diversity covering common approaches in PPI classification. F1-F5 correspond to five representative methods used in the 2012 Park and Marcotte study (Park and Marcotte 2012). In F1 (Martin, Roe, and Faulon 2005), a signature molecular descriptor (Churchwell et al. 2004; Faulon, Churchwell, and Visco 2003; Visco et al. 2002) represents each protein by the frequencies of amino acids in 3-mer combinations. F3 categorizes the 20 amino acids into seven groups according to their physicochemical properties (Shen et al. 2007); each protein sequence is then represented by the frequency of each possible 3-mer combination of these groups. F4 (Guo et al. 2008) accounts for the amino acid neighborhood context via an autocorrelation descriptor of seven physicochemical properties of each amino acid. F2 (Vert, Qiu, and Noble 2007), F5 (Park and Marcotte 2012), and F7 (Sun et al. 2017) use the same protein descriptors as in F1, F4, and F3, respectively. We introduced F6, a sequence-based domain profile method, to increase the diversity of the examined feature extraction methods. In F6, each protein is represented by its domain profile, generated by scoring the alignment of the protein amino acid sequence to the HMM profiles of 16,712 domains downloaded from Pfam in January 2018.

F1-F4 and F6 use support vector machines (SVMs) with different kernels. F6 uses the kernel in Equation [1] where (A,B) and (C,D) are two pairs of proteins. F5 utilizes a random forest classifier while F7 utilizes a stacked autoencoder (a deep learning representational learning model). Further details of F1-F5 and F7 can be found in their respective publications (Park and Marcotte 2012; Sun et al. 2017; Martin, Roe, and Faulon 2005; Vert, Qiu, and Noble 2007; Shen et al. 2007; Guo et al. 2008). We implemented the seven methods using MATLAB^R^ and paired it with LibSVM library (Chang and Lin 2011) for the SVM methods (F1-F4 and F6). 

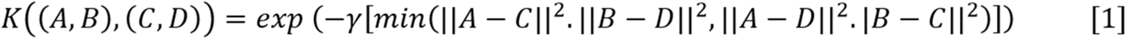

#### Benchmarking

D1-D3 were used for benchmarking the performance of the seven PPI classifiers, F1-F7. Model optimization was performed over 10-20 different combinations of the ML model hyperparameter values. Overall, we have examined 100 different models. Each was trained and tested on the 10 splits of each dataset, totaling 3,000 experiments. We did not limit benchmarking to the models with optimized hyperparameter values. The best performing models were noted for further comparisons.

#### Auditors

In AI auditing, an auxiliary ML model is designed to systematically examine bias hypotheses of an ML model of interest (main model) or its training data using the latter model input and output; a performance measure comparing the two models is defined to assess the hypotheses (Zou and Schiebinger 2018).

##### Generalizability Auditors

Two generalizability auditors, G1 and G2, were used to assess the in-network performance generalization to independent datasets before intervention (bias interrogation step) and after debiasing (bias elimination step), respectively. The main models in both auditors are the seven models optimized for D1 and D2. However, the training data for G1 is the one used for benchmarking whereas the training data for G2 is debiased first as explained in the *Debiasing Auditor* below. The test examples for the main models are subsets of D1 and D2 that satisfy the in-network performance criteria as in regular benchmarking. The auxiliary models are the same as the main models. However, the test examples for the auxiliary models in both auditors are sampled from D3 such that they satisfy the in-network test criteria for each training round. The generalizability gap was used to assess the difference in performance: the generalizability gap is the difference between the reported performance on the benchmarking test datasets (main model) and the performance on independent datasets (auxiliary model). When the gap is large, this implies that the main model does not generalize well.

##### *Feature Auditor* (A1)

The PPI classifier of interest is used as both the main model and the auxiliary model with the same hyperparameter values. In the auxiliary model, a random feature vector is constructed for each protein and used throughout the auditing experiment: each protein sequence is replaced with a random amino acid sequence before extracting the protein features. The difference in AUC of the auxiliary model to a random classifier performance (AUC ∼0.5) is used to assess the randomization efficiency.

##### *Node-degree* (A2) and *Recurrence Auditors* (A3)

The main model in both auditors is the best performing model for each benchmarking dataset (a single model per dataset). In A2, the auxiliary model is a simple (random forest) PPI classifier trained on the node degree of each protein in the positive and negative training networks (each PPI example is thus represented by a feature vector of length four). In A3, the auxiliary model is not an ML model but a scoring function that compares the summation of the node degrees of the protein pair in the positive and negative training networks. For protein pair A-B, whose positive and negative node degrees in the training data are (A^+^, B^+^) and (A^-^, B^-^), respectively, the score (interaction probability of the pair) can be described as in Equation [2]. The auxiliary models in both cases were evaluated on the 10 splits of each dataset and the quality of replication was assessed by the AUC decrease relative to the AUC obtained for the main model. In A2 and A3, there is one auditor for each dataset such that performance is compared to the best performing PPI classifier for that dataset. 

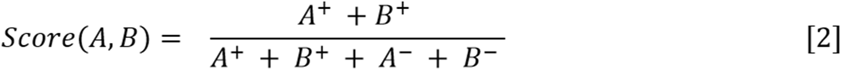

##### Debiasing Auditor

The main model is the same as in A1. For the auxiliary model, the negative examples of each data split are restricted such that each protein in the training contributes an equal number of positive and negative training examples according to the balanced sampling technique described by Yu et al. 2010, (Yu et al. 2010) which presents an unbiased alternative for random sampling. However, there was insufficient evidence to support the approach’s utility in removing bias (Ben-Hur and Noble 2006; Park and Marcotte 2011, 2012; Hamp and Rost 2015b, [a] 2015; Yu et al. 2010)). Other debiasing strategies for ML models or training data can be designed as needed.

The features of each protein were replaced by random numbers as in A1. For the asymmetric classifiers, i.e., F4, F5, and F7, which treat a pair (A,B) differently from (B,A), we accounted for interaction symmetricity (non-directionality of protein interactions) by utilizing the debiased sets prepared for the symmetric learners and representing each interaction (A,B) in the training data with the pairs (A,B) and (B,A).

Removing the representational bias was impractical for D2 as only 1,294 out of 2,181 proteins in the negative example pool are shared with the positive pool, which has 9,449 proteins. As the original negative examples were created by pairing non-co-localized proteins, we downloaded the GO localization annotation (Ashburner et al. 2000) of the proteins in D2 and split them into the following high-level co-localization groups: cytoplasm, nucleus, mitochondria, and exocytic. We constructed the negative pool by pairing all proteins that do not share a subcellular location (the same way that negative sampling was originally performed for D2) and randomly selected a subset that balances each positive training set.

Throughout the experiments, the positive and negative training example counts remain equal and the test sets remain the same as in the benchmarking data splits. The randomization efficiency is assessed as in A1.

#### Drug-target bioactivity prediction auditing

We considered five drug-target bioactivity prediction frameworks that predict whether (classification mode) and how strongly (regression mode) a drug can bind to a human protein target. All regression predictors are set up to predict the bioactivity response pKd (-log10 of the equilibrium dissociation constant Kd) while the classification models are set up to predict the binary binding status with pKd = 6.3 (corresponds to 500 nM Kd) used as the standard threshold for classification(Liu et al. 2019). AUC is used to assess classifier performances while R^2^ is used for regression models. We utilized two widely-used datasets in drug-target bioactivity research: the Metz dataset (Metz et al. 2011) and a subset of the Drug Target Commons (DTC) dataset (Tang et al. 2018), denoted here as D4 and D5, respectively. D4 and D5 consist of 107,791 and 26,634 data points measured for the bioactivity of 1,497 drugs with 172 targets and 4,210 drugs with 599 targets, respectively.

The first predictor is a classic drug-target bioactivity predictor (Cao et al. 2013) that utilizes random forest models, representing drugs with their daylight fingerprints and targets with their CTD descriptor values (Composition-Transition-Distribution standard descriptors). We re-implemented the predictor for the lack of code availability and utilized it in two modes: classification mode as F8 and regression mode as F9. The second drug-target bioactivity predictor, KronRLS (Pahikkala et al. 2015), was used in classification mode as F10 and in regression mode as F11. KronRLS represents drug and target features in a kernalized form: Smith–Waterman (SW) score for target sequences; 2D and 3D Tanimoto coefficients for the structural fingerprints of the drugs. KronRLS imputes the missing values in the drug-target all-versus-all matrix and uses the imputed values for training (but not for testing) utilizing the Kronecker RLS model (Pahikkala et al. 2013). We used the published code available for KronRLS and modified it to avoid the class imbalance problem. We changed the classification threshold and the evaluation criteria as described above. The last predictor is a recent deep learning-based classifier: DeepConv-DTI (Lee, Keum, and Nam 2019), a convolutional neural network classification model that processes target amino acid sequences directly and uses Morgan fingerprint as drug features.

The in-network performance for each framework, F8-F12, on D4 and D5 is used as the benchmarking performance (module 1) while the out-of-network performance (where both the drug and target in a test pair do not have examples of their other measurements in the training dataset) is used as the generalization performance (module 2). To remove the potential node-degree bias, we need to apply the balanced sampling discussed in the PPI prediction auditing. However, it was not feasible because the training datasets act as sparse bipartite graphs in the classification mode and have continuous output values in the regression mode. There are no distinct classes to balance the node degrees between them.

#### MHC-peptide binding prediction auditing

We considered a set of eight well known predictors of MHC class I and class II binding peptides, F13-F20, that are recently benchmarked in the 2018 Merck study (Zhao and Sher 2018): SMM-align (Nielsen, Lundegaard, and Lund 2007), Comblib (Bui et al. 2005), MHCflurry (Bui et al. 2005; O’Donnell et al. 2018), SMM^PMBEC^ (Kim et al. 2009), PickPocket (H. Zhang, Lund, and Nielsen 2009), TEPITOPE (Sturniolo et al. 1999; L. Zhang et al. 2012), NN-align (Nielsen and Lund 2009), and NetMHCpan-4 (Jurtz et al. 2017). Testing MHC-peptide binding predictors is generally performed in the out-of-network prediction mode, where the MHC allele in a test pair has examples of its binding peptides in the training set but the peptide in that test pair is novel. To assess whether these models are biased using the *Generalizability Auditor*, we compared their reported performances in their respective publications (module 1, benchmarking) with their performances on an independent dataset from the Merck study. We examined the architecture of the predictors in their respective publications and found that only PickPocket and NetMHCpan4 utilize paired-input settings. The two predictors represent the MHC alleles in terms of the amino acid sequence of their structurally identified pockets. The auditing process was stopped after module 2 as no significant bias was evident for the two paired-input models; the six other models bypass node-degree bias by design and generalize well.

## Supplementary Discussion

### Generalizability of PPI classifiers

PPI classifiers are known to have poor universal generalization; a PPI classifier trained on interactions between proteins A, B, C, D, E, and F will achieve significantly higher performance predicting new interactions between these proteins (in-network prediction) than interactions involving proteins with no examples in the training (*e.g.*, A-X, B-Y, or X-Y; out-of-network prediction) (Park and Marcotte 2012; Hamp and Rost 2015b). As a potential explanation for the high in-network performance, Park and Marcotte suggested that in-network predictions may benefit from node degree imbalances between the positive and negative training networks (Park and Marcotte 2011).

In this study, we examine whether the high in-network performance of PPI classifiers reflects true learning of protein features as Park and Marcotte suggested. If the classifiers truly learnt from protein features to make PPI predictions, they should generalize well to independent in-network test examples; the in-network performance should not change dramatically when the test PPI examples, sampled from an independent dataset, only involve proteins that are also present in the training dataset. For the purposes of testing this form of in-network generalizability, we ensured that each protein had distinct PPIs that were present in the training examples and test examples. In other words, we would only train a classifier on a subset of the known positive and negative interactions, and reserve a non-overlapping subset of the known interactions for testing. The test examples from the independent dataset were curated so that they did not include PPIs identical to those in the utilized subset of the training examples.

Indeed, our auditing framework demonstrated that classifiers do not generalize to in-network test examples from independent datasets, effectively demonstrating that the high classifier performance is not solely driven by true learning of protein features. Instead, we demonstrate that the PPI classifiers learn the node degree bias of the training examples and use this as the sole information to make predictions. The node degree of a given protein changes between datasets in contrast to the protein features of a given protein that do not change regardless of the dataset. Hence, a classifier that predominantly learns from node degrees in a training dataset is incapable of accurately predicting interactions between those proteins in an independent dataset with a different node degree bias. This illustrates the need for auditing frameworks, such as the one we designed, to systematically identify unexpected biases that impact the generalizability of classifiers.

### Node degree bias

Node degree bias drives classifier performance as follows: All interactions (e.g., A-X, B-Y) that involve either protein A or B share an identical half of their feature vectors to that of the A-B interaction. For example, if the training dataset has pairs A-B and C-D as positive and E-F and A-G as negative training examples, the A-C interaction is evaluated as being more similar to two positive training pairs and only one negative pair, resulting in a ∼⅔ probability of being a positive interaction. The classifier function can thus be described by the scoring function in Supplementary Equation [2] where the predicted score is the sum of A and B’s node degrees in the positive training network relative to the sum of their node degrees in both the positive and negative training networks. When we repeated the benchmarking of classifiers after removing node degree biases from the training datasets (balanced sampling was adopted as opposed to random sampling), classifier performance dropped significantly: the average drop in AUC was 0.19, 0.35, and 0.23 for D1, D2, and D3, respectively (**Supplementary Table 1**). This suggests that existing classifiers can be improved upon to derive more accurate PPI predictions

---

**Supplementary Figure 1:**
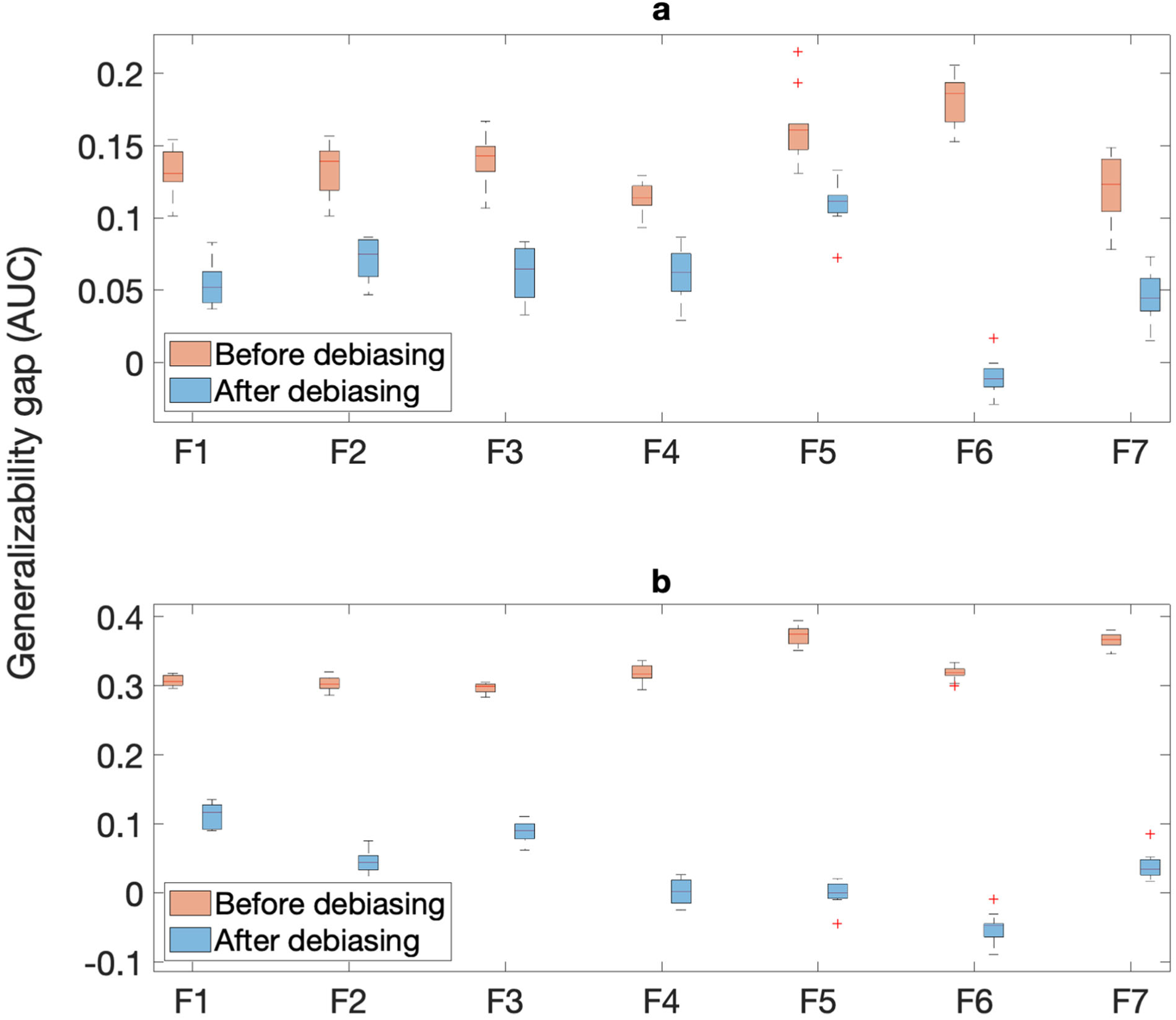
Performance gap (measured by the difference in AUC between reported/ benchmarked and independent testing performances) for the seven PPI classifiers, F1-F7, when benchmarked on datasets D1 **(a)** and D2 **(b)** and tested on dataset D3 before (pink) and after (blue) removing the node degree representational bias.

**Supplementary Table 1.** Performance based on parameter settings. Performance (measured in average AUC across 10 rounds) of frameworks F1-F7 on datasets D1-D3 using different parameter settings (P) in the following contexts: benchmarking (B), Feature Auditor (A_F_), debiased (D), and Debiasing Auditor (A_D_). For frameworks F1, F3, F4, and F6, C is the regularization parameter and *γ* is the SVM kernel coefficient. Similarly, C is the regularization parameter for F2. T and M are the number of trees and minimum leaf per node, respectively, for the random forests in F5. H and R are the hidden layer size and regularization parameter, respectively, for the autoencoder in F7.

| F | P | D1 |  |  |  | D2 |  |  |  | D3 |  |  |  |
| --- | --- | --- | --- | --- | --- | --- | --- | --- | --- | --- | --- | --- | --- |
| | | B | $A_F$ | D | $A_D$ | B | $A_F$ | D | $A_D$ | B | $A_F$ | D | $A_D$ |
| F1 | C=100, $\gamma=1e-1$ | 0.826 | 0.789 | 0.650 | 0.502 | 0.938 | 0.908 | 0.705 | 0.572 | 0.923 | 0.874 | 0.790 | 0.603 |
| | C=100, $\gamma=1e-2$ | 0.819 | 0.787 | 0.643 | 0.503 | 0.936 | 0.909 | 0.700 | 0.572 | 0.921 | 0.873 | 0.782 | 0.587 |
| | C=100, $\gamma=1e-3$ | 0.789 | 0.766 | 0.632 | 0.501 | 0.911 | 0.864 | 0.709 | 0.574 | 0.881 | 0.828 | 0.755 | 0.532 |
| | C=100, $\gamma=1e-4$ | 0.683 | 0.634 | 0.577 | 0.501 | 0.745 | 0.675 | 0.677 | 0.574 | 0.742 | 0.779 | 0.716 | 0.524 |
| | C=100, $\gamma=1e-5$ | 0.642 | 0.633 | 0.577 | 0.501 | 0.674 | 0.668 | 0.677 | 0.574 | 0.726 | 0.778 | 0.716 | 0.522 |
| | C=10, $\gamma=1e-1$ | 0.826 | 0.789 | 0.651 | 0.502 | 0.938 | 0.908 | 0.705 | 0.572 | 0.923 | 0.874 | 0.790 | 0.603 |
| | C=10, $\gamma=1e-2$ | 0.789 | 0.773 | 0.632 | 0.501 | 0.910 | 0.861 | 0.709 | 0.574 | 0.888 | 0.841 | 0.767 | 0.533 |
| | C=10, $\gamma=1e-3$ | 0.683 | 0.623 | 0.577 | 0.497 | 0.745 | 0.675 | 0.677 | 0.574 | 0.742 | 0.781 | 0.718 | 0.532 |
| | C=10, $\gamma=1e-4$ | 0.642 | 0.604 | 0.577 | 0.497 | 0.674 | 0.668 | 0.677 | 0.574 | 0.726 | 0.779 | 0.716 | 0.524 |
| | C=10, $\gamma=1e-5$ | 0.642 | 0.604 | 0.577 | 0.497 | 0.674 | 0.668 | 0.677 | 0.574 | 0.726 | 0.778 | 0.716 | 0.522 |
| | C=1, $\gamma=1e-1$ | 0.781 | 0.760 | 0.630 | 0.497 | 0.902 | 0.833 | 0.707 | 0.574 | 0.881 | 0.834 | 0.794 | 0.624 |
| | C=1, $\gamma=1e-2$ | 0.682 | 0.621 | 0.577 | 0.497 | 0.742 | 0.674 | 0.678 | 0.574 | 0.745 | 0.794 | 0.728 | 0.532 |
| | C=1, $\gamma=1e-3$ | 0.642 | 0.604 | 0.577 | 0.497 | 0.674 | 0.668 | 0.677 | 0.574 | 0.726 | 0.781 | 0.718 | 0.532 |
| | C=1, $\gamma=1e-4$ | 0.642 | 0.604 | 0.577 | 0.497 | 0.674 | 0.668 | 0.677 | 0.574 | 0.726 | 0.779 | 0.716 | 0.524 |
| | C=1, $\gamma=1e-5$ | 0.642 | 0.604 | 0.580 | 0.497 | 0.674 | 0.668 | 0.677 | 0.574 | 0.726 | 0.778 | 0.716 | 0.522 |
| F2 | C=1e3 | 0.811 | 0.799 | 0.586 | 0.512 | 0.953 | 0.969 | 0.602 | 0.578 | 0.898 | 0.850 | 0.636 | 0.310 |
|  | C=5e2 | 0.811 | 0.799 | 0.586 | 0.512 | 0.953 | 0.969 | 0.602 | 0.578 | 0.898 | 0.850 | 0.636 | 0.310 |
|  | C=1e2 | 0.811 | 0.799 | 0.586 | 0.512 | 0.953 | 0.969 | 0.602 | 0.578 | 0.898 | 0.850 | 0.636 | 0.310 |
|  | C=5e1 | 0.811 | 0.799 | 0.586 | 0.512 | 0.953 | 0.969 | 0.602 | 0.578 | 0.898 | 0.850 | 0.636 | 0.310 |
|  | C=1e1 | 0.811 | 0.799 | 0.586 | 0.512 | 0.953 | 0.969 | 0.602 | 0.578 | 0.898 | 0.850 | 0.636 | 0.310 |
|  | C=5e0 | 0.811 | 0.799 | 0.586 | 0.512 | 0.953 | 0.969 | 0.602 | 0.578 | 0.898 | 0.850 | 0.636 | 0.310 |
|  | C=1e0 | 0.811 | 0.799 | 0.586 | 0.512 | 0.953 | 0.969 | 0.602 | 0.578 | 0.898 | 0.850 | 0.636 | 0.310 |
|  | C=5e-1 | 0.811 | 0.799 | 0.586 | 0.512 | 0.953 | 0.969 | 0.602 | 0.578 | 0.898 | 0.850 | 0.636 | 0.310 |
|  | C=1e-1 | 0.811 | 0.799 | 0.586 | 0.512 | 0.953 | 0.969 | 0.602 | 0.578 | 0.898 | 0.850 | 0.636 | 0.310 |
|  | C=5e-2 | 0.811 | 0.799 | 0.586 | 0.512 | 0.953 | 0.969 | 0.602 | 0.578 | 0.898 | 0.850 | 0.635 | 0.310 |
| F3 | C=100, $\gamma=1e-1$ | 0.681 | 0.642 | 0.550 | 0.495 | 0.901 | 0.913 | 0.612 | 0.572 | 0.768 | 0.728 | 0.647 | 0.508 |
| | C=100, $\gamma=1e-2$ | 0.577 | 0.597 | 0.536 | 0.536 | 0.669 | 0.797 | 0.574 | 0.530 | 0.660 | 0.670 | 0.620 | 0.562 |
| | C=100, $\gamma=1e-3$ | 0.541 | 0.580 | 0.509 | 0.502 | 0.619 | 0.645 | 0.540 | 0.486 | 0.633 | 0.658 | 0.543 | 0.514 |
| | C=100, $\gamma=1e-4$ | 0.517 | 0.547 | 0.507 | 0.506 | 0.571 | 0.603 | 0.544 | 0.495 | 0.557 | 0.605 | 0.521 | 0.492 |
| | C=100, $\gamma=1e-5$ | 0.514 | 0.532 | 0.506 | 0.505 | 0.567 | 0.581 | 0.543 | 0.495 | 0.546 | 0.572 | 0.513 | 0.489 |
| | C=10, $\gamma=1e-1$ | 0.686 | 0.642 | 0.552 | 0.495 | 0.901 | 0.913 | 0.615 | 0.572 | 0.768 | 0.672 | 0.648 | 0.508 |
| | C=100, $\gamma=1e-2$ | 0.598 | 0.647 | 0.545 | 0.527 | 0.671 | 0.901 | 0.555 | 0.554 | 0.681 | 0.699 | 0.615 | 0.553 |
| | C=10, $\gamma=1e-3$ | 0.541 | 0.586 | 0.509 | 0.495 | 0.624 | 0.654 | 0.540 | 0.490 | 0.634 | 0.671 | 0.534 | 0.514 |
| | C=10, $\gamma=1e-4$ | 0.517 | 0.547 | 0.507 | 0.506 | 0.572 | 0.604 | 0.544 | 0.495 | 0.558 | 0.607 | 0.521 | 0.494 |
| | C=10, $\gamma=1e-5$ | 0.515 | 0.533 | 0.506 | 0.506 | 0.569 | 0.583 | 0.543 | 0.493 | 0.551 | 0.579 | 0.513 | 0.520 |
| | C=1, $\gamma=1e-1$ | 0.700 | 0.642 | 0.555 | 0.495 | 0.904 | 0.913 | 0.619 | 0.572 | 0.769 | 0.672 | 0.646 | 0.508 |
| | C=1, $\gamma=1e-2$ | 0.587 | 0.716 | 0.541 | 0.481 | 0.697 | 0.942 | 0.544 | 0.551 | 0.819 | 0.770 | 0.643 | 0.510 |
| | C=1, $\gamma=1e-3$ | 0.542 | 0.590 | 0.509 | 0.499 | 0.630 | 0.670 | 0.540 | 0.485 | 0.645 | 0.708 | 0.544 | 0.533 |
| | C=1, $\gamma=1e-4$ | 0.518 | 0.549 | 0.505 | 0.507 | 0.575 | 0.611 | 0.544 | 0.493 | 0.566 | 0.628 | 0.522 | 0.516 |
| | C=1, $\gamma=1e-5$ | 0.520 | 0.542 | 0.506 | 0.499 | 0.611 | 0.532 | 0.522 | 0.493 | 0.688 | 0.712 | 0.538 | 0.508 |
| F4 | C=100, $\gamma=1e-1$ | 0.664 | 0.671 | 0.527 | 0.502 | 0.899 | 0.914 | 0.588 | 0.573 | 0.797 | 0.758 | 0.629 | 0.525 |
| | C=100, $\gamma=1e-2$ | 0.680 | 0.697 | 0.543 | 0.511 | 0.893 | 0.911 | 0.570 | 0.579 | 0.816 | 0.800 | 0.628 | 0.542 |
| | C=100, $\gamma=1e-3$ | 0.700 | 0.713 | 0.550 | 0.511 | 0.873 | 0.883 | 0.550 | 0.545 | 0.827 | 0.828 | 0.591 | 0.552 |
| | C=100, $\gamma=1e-4$ | 0.647 | 0.650 | 0.531 | 0.496 | 0.777 | 0.759 | 0.531 | 0.506 | 0.760 | 0.766 | 0.563 | 0.525 |
| | C=100, $\gamma=1e-5$ | 0.593 | 0.602 | 0.505 | 0.500 | 0.681 | 0.685 | 0.524 | 0.496 | 0.669 | 0.722 | 0.529 | 0.508 |
| | C=10, $\gamma=1e-1$ | 0.667 | 0.677 | 0.531 | 0.502 | 0.899 | 0.915 | 0.587 | 0.573 | 0.797 | 0.758 | 0.629 | 0.525 |
| | C=100, $\gamma=1e-2$ | 0.705 | 0.718 | 0.557 | 0.510 | 0.912 | 0.926 | 0.581 | 0.579 | 0.837 | 0.824 | 0.634 | 0.542 |
| | C=10, $\gamma=1e-3$ | 0.683 | 0.688 | 0.546 | 0.497 | 0.836 | 0.843 | 0.549 | 0.520 | 0.809 | 0.805 | 0.578 | 0.522 |
| | C=10, $\gamma=1e-4$ | 0.611 | 0.619 | 0.522 | 0.495 | 0.707 | 0.708 | 0.540 | 0.504 | 0.703 | 0.737 | 0.556 | 0.509 |
| | C=10, $\gamma=1e-5$ | 0.567 | 0.593 | 0.515 | 0.500 | 0.625 | 0.665 | 0.528 | 0.496 | 0.618 | 0.684 | 0.536 | 0.508 |
| | C=1, $\gamma=1e-1$ | 0.679 | 0.692 | 0.545 | 0.502 | 0.898 | 0.921 | 0.590 | 0.573 | 0.798 | 0.771 | 0.629 | 0.524 |
| | C=1, $\gamma=1e-2$ | 0.701 | 0.710 | 0.570 | 0.497 | 0.879 | 0.899 | 0.574 | 0.573 | 0.826 | 0.821 | 0.611 | 0.532 |
| | C=1, $\gamma=1e-3$ | 0.640 | 0.652 | 0.550 | 0.499 | 0.755 | 0.768 | 0.555 | 0.504 | 0.747 | 0.769 | 0.587 | 0.511 |
| | C=1, $\gamma=1e-4$ | 0.575 | 0.605 | 0.522 | 0.500 | 0.639 | 0.668 | 0.545 | 0.495 | 0.634 | 0.690 | 0.555 | 0.508 |
| | C=1, $\gamma=1e-5$ | 0.543 | 0.564 | 0.515 | 0.500 | 0.578 | 0.576 | 0.529 | 0.496 | 0.581 | 0.580 | 0.536 | 0.508 |
| F5 | T=1e1, M=1 | 0.700 | 0.703 | 0.522 | 0.507 | 0.974 | 0.989 | 0.568 | 0.567 | 0.875 | 0.861 | 0.600 | 0.536 |
|  | T=1e1, M=5 | 0.719 | 0.727 | 0.516 | 0.504 | 0.971 | 0.990 | 0.563 | 0.561 | 0.887 | 0.875 | 0.600 | 0.525 |
|  | T=1e1, M=10 | 0.726 | 0.726 | 0.521 | 0.504 | 0.968 | 0.991 | 0.563 | 0.548 | 0.884 | 0.877 | 0.583 | 0.523 |
|  | T=1e1, M=50 | 0.728 | 0.736 | 0.528 | 0.497 | 0.929 | 0.991 | 0.542 | 0.529 | 0.873 | 0.871 | 0.560 | 0.518 |
|  | T=5e1, M=1 | 0.760 | 0.766 | 0.548 | 0.514 | 0.985 | 0.992 | 0.592 | 0.576 | 0.902 | 0.888 | 0.655 | 0.557 |
|  | T=5e1, M=5 | 0.768 | 0.775 | 0.547 | 0.504 | 0.983 | 0.993 | 0.587 | 0.573 | 0.902 | 0.892 | 0.649 | 0.544 |
|  | T=5e1, M=10 | 0.772 | 0.777 | 0.547 | 0.509 | 0.981 | 0.993 | 0.586 | 0.572 | 0.900 | 0.893 | 0.631 | 0.538 |
|  | T=5e1, M=50 | 0.764 | 0.775 | 0.545 | 0.506 | 0.963 | 0.992 | 0.567 | 0.556 | 0.887 | 0.885 | 0.603 | 0.535 |
|  | T=1e2, M=1 | 0.774 | 0.777 | 0.563 | 0.527 | 0.986 | 0.993 | 0.595 | 0.576 | 0.905 | 0.892 | 0.675 | 0.559 |
|  | T=1e2, M=5 | 0.778 | 0.784 | 0.558 | 0.513 | 0.984 | 0.993 | 0.597 | 0.578 | 0.905 | 0.894 | 0.656 | 0.564 |
|  | T=1e2, M=10 | 0.779 | 0.786 | 0.561 | 0.503 | 0.983 | 0.993 | 0.589 | 0.572 | 0.904 | 0.896 | 0.653 | 0.557 |
|  | T=1e2, M=50 | 0.771 | 0.784 | 0.554 | 0.504 | 0.966 | 0.992 | 0.583 | 0.565 | 0.889 | 0.888 | 0.615 | 0.534 |
|  | T=5e2, M=1 | 0.784 | 0.788 | 0.584 | 0.531 | 0.986 | 0.993 | 0.603 | 0.581 | 0.908 | 0.895 | 0.697 | 0.588 |
|  | T=5e2, M=5 | 0.787 | 0.793 | 0.578 | 0.521 | 0.985 | 0.989 | 0.603 | 0.580 | 0.907 | 0.897 | 0.686 | 0.570 |
|  | T=5e2, M=10 | 0.783 | 0.794 | 0.572 | 0.512 | 0.984 | 0.988 | 0.605 | 0.576 | 0.905 | 0.897 | 0.676 | 0.561 |
|  | T=5e2, M=50 | 0.783 | 0.790 | 0.569 | 0.508 | 0.968 | 0.984 | 0.598 | 0.572 | 0.891 | 0.891 | 0.643 | 0.539 |
|  | T=1e3, M=1 | 0.782 | 0.790 | 0.591 | 0.532 | 0.986 | 0.990 | 0.604 | 0.580 | 0.908 | 0.896 | 0.700 | 0.590 |
|  | T=1e3, M=5 | 0.788 | 0.794 | 0.581 | 0.522 | 0.985 | 0.989 | 0.606 | 0.578 | 0.907 | 0.898 | 0.693 | 0.571 |
|  | T=1e3, M=10 | 0.787 | 0.794 | 0.576 | 0.515 | 0.984 | 0.988 | 0.603 | 0.580 | 0.906 | 0.898 | 0.684 | 0.564 |
|  | T=1e3, M=50 | 0.776 | 0.791 | 0.572 | 0.506 | 0.969 | 0.984 | 0.601 | 0.576 | 0.891 | 0.891 | 0.655 | 0.547 |
| F6 | C=100, $\gamma=1e-1$ | 0.798 | 0.799 | 0.500 | 0.500 | 0.885 | 0.886 | 0.383 | 0.427 | 0.823 | 0.828 | 0.436 | 0.435 |
| | C=100, $\gamma=1e-2$ | 0.795 | 0.798 | 0.499 | 0.534 | 0.898 | 0.886 | 0.443 | 0.396 | 0.825 | 0.828 | 0.424 | 0.435 |
| | C=100, $\gamma=1e-3$ | 0.803 | 0.801 | 0.534 | 0.419 | 0.912 | 0.900 | 0.481 | 0.415 | 0.809 | 0.835 | 0.417 | 0.413 |
| | C=100, $\gamma=1e-4$ | 0.593 | 0.800 | 0.499 | 0.498 | 0.669 | 0.910 | 0.488 | 0.429 | 0.626 | 0.809 | 0.480 | 0.488 |
| | C=100, $\gamma=1e-5$ | 0.541 | 0.798 | 0.499 | 0.497 | 0.597 | 0.696 | 0.497 | 0.428 | 0.586 | 0.726 | 0.497 | 0.485 |
| | C=10, $\gamma=1e-1$ | 0.801 | 0.801 | 0.500 | 0.500 | 0.959 | 0.962 | 0.383 | 0.427 | 0.824 | 0.829 | 0.438 | 0.434 |
| | C=100, $\gamma=1e-2$ | 0.801 | 0.801 | 0.499 | 0.534 | 0.961 | 0.962 | 0.444 | 0.396 | 0.830 | 0.829 | 0.424 | 0.434 |
| | C=10, $\gamma=1e-3$ | 0.806 | 0.803 | 0.537 | 0.492 | 0.956 | 0.964 | 0.482 | 0.418 | 0.819 | 0.836 | 0.411 | 0.447 |
| | C=10, $\gamma=1e-4$ | 0.594 | 0.804 | 0.499 | 0.498 | 0.674 | 0.925 | 0.488 | 0.429 | 0.631 | 0.822 | 0.480 | 0.463 |
| | C=10, $\gamma=1e-5$ | 0.541 | 0.811 | 0.500 | 0.498 | 0.603 | 0.717 | 0.497 | 0.429 | 0.586 | 0.744 | 0.498 | 0.463 |
| | C=1, $\gamma=1e-1$ | 0.824 | 0.824 | 0.500 | 0.500 | 0.981 | 0.981 | 0.383 | 0.427 | 0.853 | 0.856 | 0.432 | 0.427 |
| | C=1, $\gamma=1e-2$ | 0.823 | 0.824 | 0.499 | 0.534 | 0.981 | 0.981 | 0.443 | 0.396 | 0.852 | 0.856 | 0.454 | 0.427 |
| | C=1, $\gamma=1e-3$ | 0.825 | 0.824 | 0.536 | 0.430 | 0.979 | 0.982 | 0.499 | 0.408 | 0.850 | 0.855 | 0.445 | 0.398 |
| | C=1, $\gamma=1e-4$ | 0.598 | 0.825 | 0.499 | 0.498 | 0.669 | 0.960 | 0.489 | 0.429 | 0.644 | 0.854 | 0.479 | 0.456 |
| | C=1, $\gamma=1e-5$ | 0.543 | 0.834 | 0.500 | 0.498 | 0.611 | 0.753 | 0.498 | 0.429 | 0.611 | 0.850 | 0.485 | 0.456 |
| F7 | H=100, R=1e-3 | 0.734 | 0.715 | 0.571 | 0.500 | 0.938 | 0.914 | 0.583 | 0.500 | 0.873 | 0.843 | 0.641 | 0.530 |
|  | H=100, R=5e-3 | 0.711 | 0.682 | 0.555 | 0.500 | 0.898 | 0.754 | 0.547 | 0.500 | 0.843 | 0.803 | 0.610 | 0.500 |
|  | H=100, R=1e-2 | 0.668 | 0.669 | 0.522 | 0.500 | 0.876 | 0.753 | 0.555 | 0.500 | 0.824 | 0.798 | 0.500 | 0.500 |
|  | H=100, R=5e-2 | 0.666 | 0.623 | 0.500 | 0.500 | 0.812 | 0.713 | 0.500 | 0.500 | 0.822 | 0.763 | 0.500 | 0.500 |
|  | H=100, R=1e-1 | 0.649 | 0.626 | 0.500 | 0.500 | 0.846 | 0.735 | 0.500 | 0.500 | 0.822 | 0.791 | 0.500 | 0.500 |
|  | H=400, R=1e-3 | 0.733 | 0.716 | 0.570 | 0.502 | 0.948 | 0.925 | 0.593 | 0.505 | 0.875 | 0.846 | 0.647 | 0.521 |
|  | H=400, R=5e-3 | 0.703 | 0.661 | 0.551 | 0.500 | 0.895 | 0.765 | 0.555 | 0.500 | 0.846 | 0.792 | 0.596 | 0.500 |
|  | H=400, R=1e-2 | 0.674 | 0.656 | 0.531 | 0.500 | 0.862 | 0.739 | 0.575 | 0.500 | 0.823 | 0.796 | 0.500 | 0.500 |
|  | H=400, R=5e-2 | 0.668 | 0.608 | 0.500 | 0.500 | 0.845 | 0.735 | 0.500 | 0.500 | 0.792 | 0.791 | 0.500 | 0.500 |
|  | H=400, R=1e-1 | 0.600 | 0.638 | 0.500 | 0.500 | 0.846 | 0.712 | 0.500 | 0.500 | 0.759 | 0.793 | 0.500 | 0.500 |

**Supplementary Table 2.**
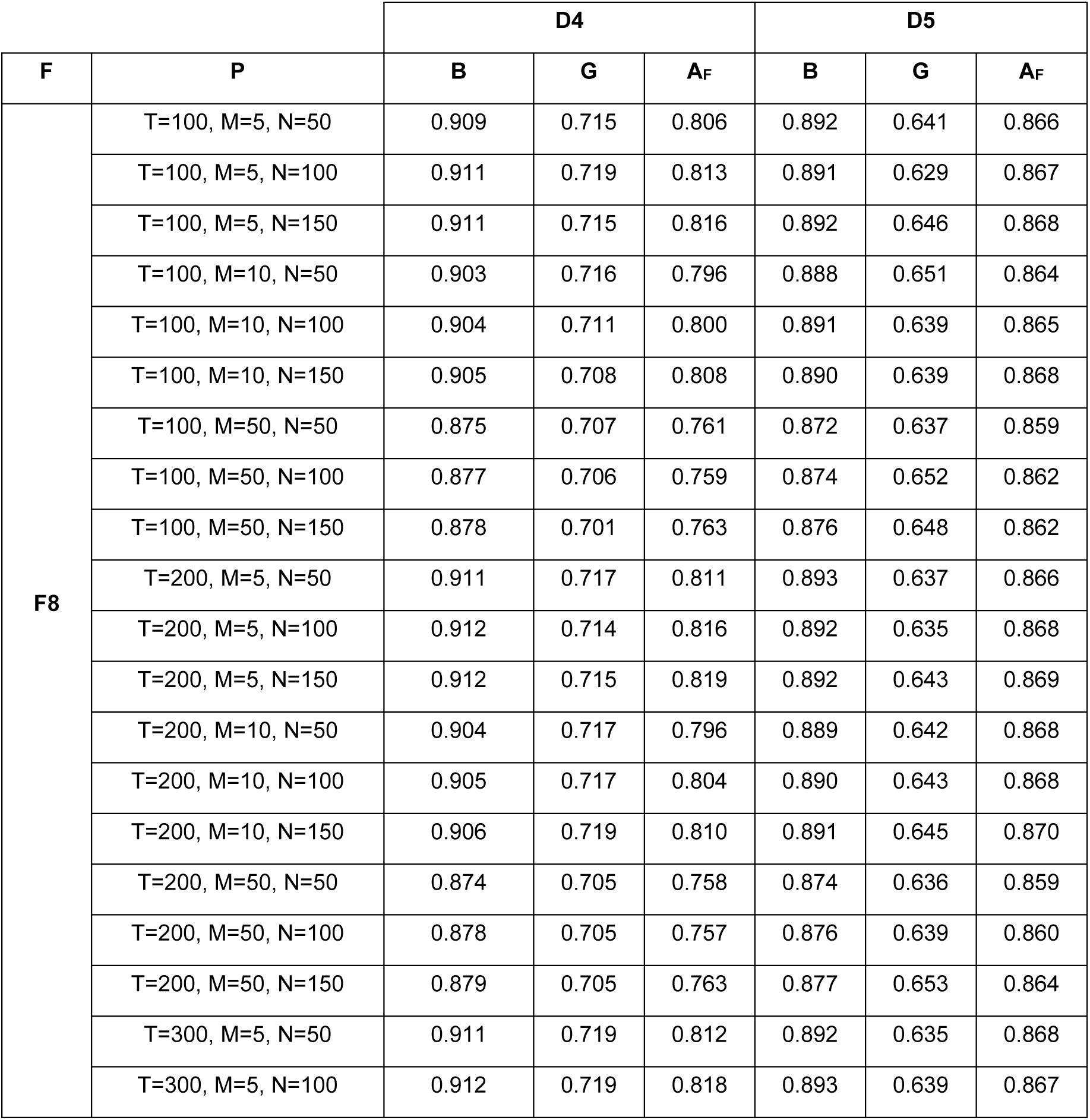
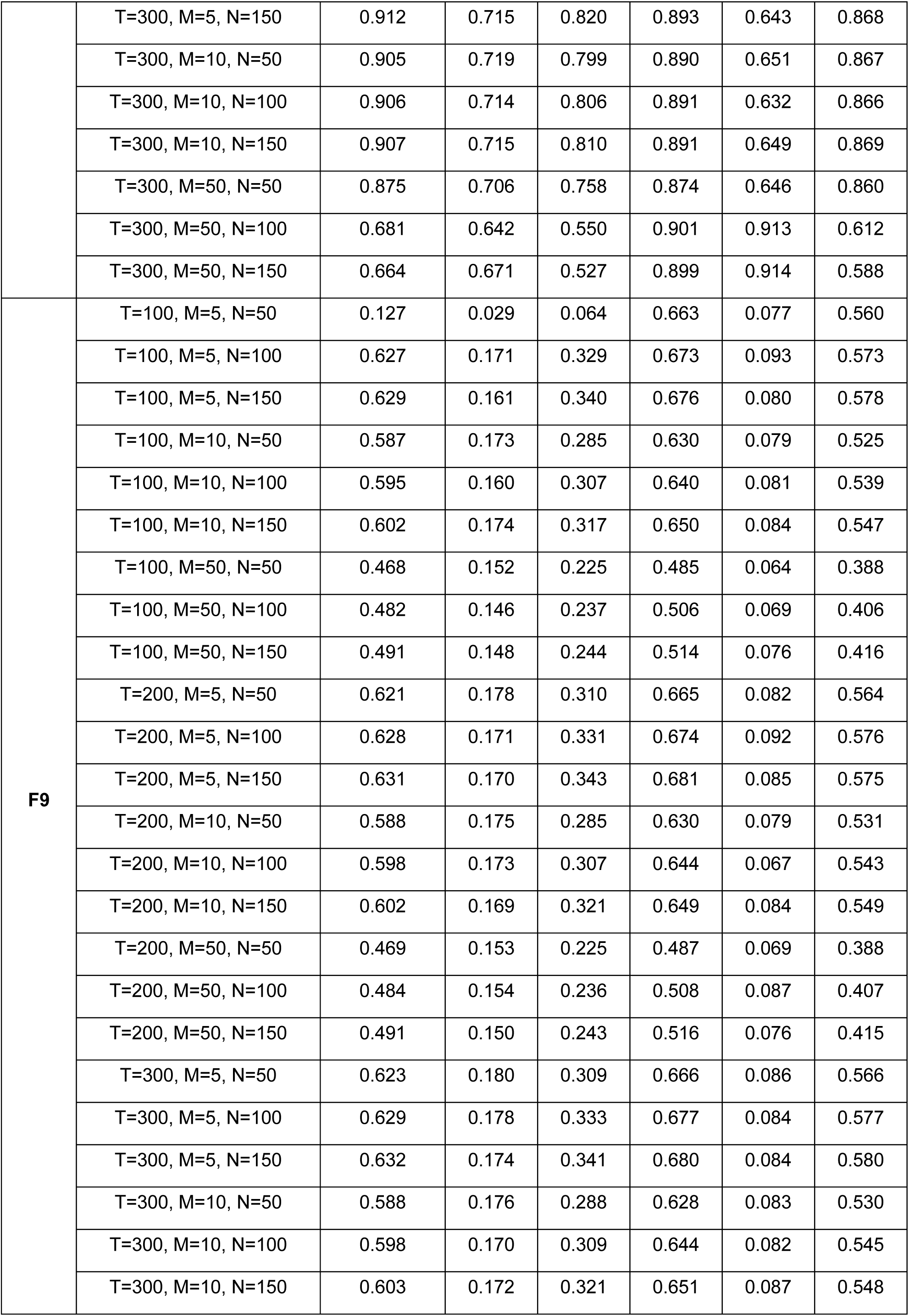
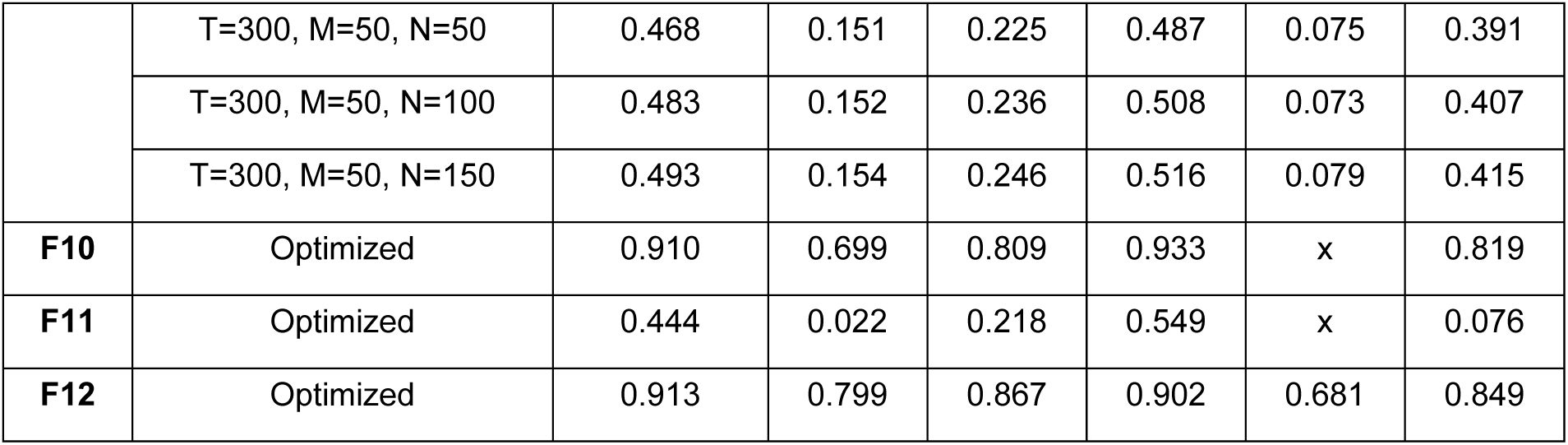
Performance based on parameter settings F8-12. Performance (measured in average AUC for the classification frameworks F8, F10, and F12, and in average R-squared for the regression frameworks F9 and F11) on datasets D4 and D5 using different parameter settings for F8 and F9, and the optimized parameters for F10, F11, and F12, in the following contexts: benchmarking (B), generalization (G) and Feature Auditor (A_F_). For F8 and F9, T is the number of trees, M is the minimum leaf per node, and N is the number of predictors per node for the random forests. The x values indicate where a model fails to operate.

| F | P | D4 |  |  | D5 |  |  |
| --- | --- | --- | --- | --- | --- | --- | --- |
| | | B | G | $A_F$ | B | G | $A_F$ |
| F8 | T=100, M=5, N=50 | 0.909 | 0.715 | 0.806 | 0.892 | 0.641 | 0.866 |
|  | T=100, M=5, N=100 | 0.911 | 0.719 | 0.813 | 0.891 | 0.629 | 0.867 |
|  | T=100, M=5, N=150 | 0.911 | 0.715 | 0.816 | 0.892 | 0.646 | 0.868 |
|  | T=100, M=10, N=50 | 0.903 | 0.716 | 0.796 | 0.888 | 0.651 | 0.864 |
|  | T=100, M=10, N=100 | 0.904 | 0.711 | 0.800 | 0.891 | 0.639 | 0.865 |
|  | T=100, M=10, N=150 | 0.905 | 0.708 | 0.808 | 0.890 | 0.639 | 0.868 |
|  | T=100, M=50, N=50 | 0.875 | 0.707 | 0.761 | 0.872 | 0.637 | 0.859 |
|  | T=100, M=50, N=100 | 0.877 | 0.706 | 0.759 | 0.874 | 0.652 | 0.862 |
|  | T=100, M=50, N=150 | 0.878 | 0.701 | 0.763 | 0.876 | 0.648 | 0.862 |
|  | T=200, M=5, N=50 | 0.911 | 0.717 | 0.811 | 0.893 | 0.637 | 0.866 |
|  | T=200, M=5, N=100 | 0.912 | 0.714 | 0.816 | 0.892 | 0.635 | 0.868 |
|  | T=200, M=5, N=150 | 0.912 | 0.715 | 0.819 | 0.892 | 0.643 | 0.869 |
|  | T=200, M=10, N=50 | 0.904 | 0.717 | 0.796 | 0.889 | 0.642 | 0.868 |
|  | T=200, M=10, N=100 | 0.905 | 0.717 | 0.804 | 0.890 | 0.643 | 0.868 |
|  | T=200, M=10, N=150 | 0.906 | 0.719 | 0.810 | 0.891 | 0.645 | 0.870 |
|  | T=200, M=50, N=50 | 0.874 | 0.705 | 0.758 | 0.874 | 0.636 | 0.859 |
|  | T=200, M=50, N=100 | 0.878 | 0.705 | 0.757 | 0.876 | 0.639 | 0.860 |
|  | T=200, M=50, N=150 | 0.879 | 0.705 | 0.763 | 0.877 | 0.653 | 0.864 |
|  | T=300, M=5, N=50 | 0.911 | 0.719 | 0.812 | 0.892 | 0.635 | 0.868 |
|  | T=300, M=5, N=100 | 0.912 | 0.719 | 0.818 | 0.893 | 0.639 | 0.867 |
|  | T=300, M=5, N=150 | 0.912 | 0.715 | 0.820 | 0.893 | 0.643 | 0.868 |
|  | T=300, M=10, N=50 | 0.905 | 0.719 | 0.799 | 0.890 | 0.651 | 0.867 |
|  | T=300, M=10, N=100 | 0.906 | 0.714 | 0.806 | 0.891 | 0.632 | 0.866 |
|  | T=300, M=10, N=150 | 0.907 | 0.715 | 0.810 | 0.891 | 0.649 | 0.869 |
|  | T=300, M=50, N=50 | 0.875 | 0.706 | 0.758 | 0.874 | 0.646 | 0.860 |
|  | T=300, M=50, N=100 | 0.681 | 0.642 | 0.550 | 0.901 | 0.913 | 0.612 |
|  | T=300, M=50, N=150 | 0.664 | 0.671 | 0.527 | 0.899 | 0.914 | 0.588 |
| **F9** | T=100, M=5, N=50 | 0.127 | 0.029 | 0.064 | 0.663 | 0.077 | 0.560 |
|  | T=100, M=5, N=100 | 0.627 | 0.171 | 0.329 | 0.673 | 0.093 | 0.573 |
|  | T=100, M=5, N=150 | 0.629 | 0.161 | 0.340 | 0.676 | 0.080 | 0.578 |
|  | T=100, M=10, N=50 | 0.587 | 0.173 | 0.285 | 0.630 | 0.079 | 0.525 |
|  | T=100, M=10, N=100 | 0.595 | 0.160 | 0.307 | 0.640 | 0.081 | 0.539 |
|  | T=100, M=10, N=150 | 0.602 | 0.174 | 0.317 | 0.650 | 0.084 | 0.547 |
|  | T=100, M=50, N=50 | 0.468 | 0.152 | 0.225 | 0.485 | 0.064 | 0.388 |
|  | T=100, M=50, N=100 | 0.482 | 0.146 | 0.237 | 0.506 | 0.069 | 0.406 |
|  | T=100, M=50, N=150 | 0.491 | 0.148 | 0.244 | 0.514 | 0.076 | 0.416 |
|  | T=200, M=5, N=50 | 0.621 | 0.178 | 0.310 | 0.665 | 0.082 | 0.564 |
|  | T=200, M=5, N=100 | 0.628 | 0.171 | 0.331 | 0.674 | 0.092 | 0.576 |
|  | T=200, M=5, N=150 | 0.631 | 0.170 | 0.343 | 0.681 | 0.085 | 0.575 |
|  | T=200, M=10, N=50 | 0.588 | 0.175 | 0.285 | 0.630 | 0.079 | 0.531 |
|  | T=200, M=10, N=100 | 0.598 | 0.173 | 0.307 | 0.644 | 0.067 | 0.543 |
|  | T=200, M=10, N=150 | 0.602 | 0.169 | 0.321 | 0.649 | 0.084 | 0.549 |
|  | T=200, M=50, N=50 | 0.469 | 0.153 | 0.225 | 0.487 | 0.069 | 0.388 |
|  | T=200, M=50, N=100 | 0.484 | 0.154 | 0.236 | 0.508 | 0.087 | 0.407 |
|  | T=200, M=50, N=150 | 0.491 | 0.150 | 0.243 | 0.516 | 0.076 | 0.415 |
|  | T=300, M=5, N=50 | 0.623 | 0.180 | 0.309 | 0.666 | 0.086 | 0.566 |
|  | T=300, M=5, N=100 | 0.629 | 0.178 | 0.333 | 0.677 | 0.084 | 0.577 |
|  | T=300, M=5, N=150 | 0.632 | 0.174 | 0.341 | 0.680 | 0.084 | 0.580 |
|  | T=300, M=10, N=50 | 0.588 | 0.176 | 0.288 | 0.628 | 0.083 | 0.530 |
|  | T=300, M=10, N=100 | 0.598 | 0.170 | 0.309 | 0.644 | 0.082 | 0.545 |
|  | T=300, M=10, N=150 | 0.603 | 0.172 | 0.321 | 0.651 | 0.087 | 0.548 |
|  | T=300, M=50, N=50 | 0.468 | 0.151 | 0.225 | 0.487 | 0.075 | 0.391 |
|  | T=300, M=50, N=100 | 0.483 | 0.152 | 0.236 | 0.508 | 0.073 | 0.407 |
|  | T=300, M=50, N=150 | 0.493 | 0.154 | 0.246 | 0.516 | 0.079 | 0.415 |
| **F10** | Optimized | 0.910 | 0.699 | 0.809 | 0.933 | x | 0.819 |
| **F11** | Optimized | 0.444 | 0.022 | 0.218 | 0.549 | x | 0.076 |
| **F12** | Optimized | 0.913 | 0.799 | 0.867 | 0.902 | 0.681 | 0.849 |

